# Developing bio-technical reliability among future researchers and industrial work-force in developing countries: a success story

**DOI:** 10.1101/2020.02.03.919209

**Authors:** Kshitish K Acharya

## Abstract

A common problem in the biotech sector of developing countries is that a large portion of students attain a poor conceptual understanding of the basic theory and/or lack proficiency in basic laboratory skills even as they complete higher studies such as a master’s degree. A small scale solution was developed in the form of a unique post-graduate diploma program that imparted reliable skills in students with good theoretical knowledge. The course of 6 to 8 months duration, with an optional 6 months internship, was successfully conducted for 8 batches. Most students of this course inculcated the right laboratory practices. This was evident by safe operations, and precise as well as accurate results of quantitative experiments conducted by them. They learned to work independently as well as in teams. They could create and follow standard operating procedures, and contribute to general laboratory maintenance. Every student also designed, prepared and conducted routine molecular biology experiments. The following aspects are suggested for successfully dealing with the patchy and varied capacities of life science students during higher (post bachelor’s) studies: a) careful selection of students, b) training with prioritized objectives, c) attention to basic lab-practices, d) opportunities to self-learn, e) structured group discussions, f) teacher(s) with genuine interest and passion, g) reasonable infrastructure, and h) maintaining a good student-instructor ratio. The course objectives, structure, teaching methods, and experiences are presented here along with some of the relevant data, including statistics about improvement in the precision and accuracy in experimental results by students. Limitations in this course and a general critical perspective of the routine higher education have also been discussed briefly.

## Introduction

With time, the education requirements within the life sciences domains have been changing (1). Developing countries such as India seem to be keeping up to some extent. There has been significant growth in the Indian Biotech sector in recent years (2, 3) and, like elsewhere (4), the job market has diversified. Correspondingly, the number of job-openings increased, but this was accompanied by a disproportionately larger surge in the number of bachelor’s and master’s graduates in the life sciences subjects (5). Only an average of about 25% of the out-going students from most education-centers seem to find jobs after their graduation – an indication by certain reports (e.g., 6) strongly supported by personal observations made over 15 years via personal interactions with academicians and students from various parts of India as well as occasional informal surveys on social media. The unfortunate situation of poor employability of young biologists, probably common to a few other developing countries, stems from various limitations of the prevailing education systems (7–10). The majority of post-bachelors-degree students from life sciences streams such as biotechnology, microbiology, biochemistry, zoology, agriculture, and botany, possess patchy capacities in terms of safe and reproducible execution of routine basic laboratory procedures. Such an incompleteness in diverse types of initial skill sets poses a huge obstacle while training the students in advanced biotechnical laboratory techniques. The irregular capacities of students may also be key causes for eventual limitations in the overall employability of many youngsters, even after a master’s degree. Overall, the following major limitations exist among most of the current job-seekers from various life science streams within India: a) poor understanding of the fundamental concepts in the core domains such as cell and molecular biology, and b) poor dependability in terms of laboratory experimental tasks. I had an opportunity to attempt and find a solution to the current problem in human resource development in the biotech sector, though on a very small scale, in the form of a post-graduate Laboratory Course in Bio-Techniques, the LCBT. I designed and conducted this course of 6 to 8 months, with an option for additional 6 months’ internship. The strategy involved careful attention to specific lacunae, time-bound prioritization of objectives and a unique combination of methods of training.

The purpose of this new program was to contribute to the technical work-force of the biotech industry as well as prepare students for academic research involving experiments in cell and molecular biology. Students with sound theoretical knowledge in vital biochemistry, cell & molecular biology were carefully selected and practical training was imparted. About 1 in 5 applicants were offered admission to the program (see supplementary notes for details of selection). The students recruited for the course still had varying degrees of initial capacities in terms of safe and correct execution of commonly required laboratory procedures. But it was possible to eventually impart the required awareness and inculcate the right laboratory practices among students before training them in advanced molecular biology techniques. We were able to attain a placement record of about 90% across 8 batches.

## Methods

Observations during the 12 years of experience preceding 2003, with industry and academia (India and abroad), helped me identify the need for such a program. Interactions with other industrial and academic experts in the domain helped the consolidation of this understanding as well as a broad plan. The first-of-its-kind program in the country was then conceptualized. The curriculum designed and implemented successfully at the Institute of Bioinformatics and Applied Biotechnology (IBAB). A combination of the following measures comprised the novel strategy adopted to achieve the training goals:

a) Careful selection of students with a sound conceptual understanding of fundamentals in the core domains [see supplementary notes]

b) Prioritized specific objectives with defined time limits. There were broadly two phases of the course in terms of the objectives as indicated in the text-boxes (box 1 & 2).

### BOX 1. Phase I objectives, for first 60 days

Mainly to ensure that ‘all’ students have acquired the following knowledge, and gained the habits and/or confidence in the listed operations.

- Awareness of hazards and a persistent attention to safety requirements and preventive steps with continuous monitoring of protocol deviations and any potential or actual accidents.
- A good understanding of the extent of variations in the results and reasons for these variations, via repeated individual performances of quantitative experiments & analysis of results.
- Laboratory mathematics and independent solution preparations.
- Optimal use of basic computational tools such as word/writer, spreadsheets and slide-/presentation-makers.
- Searching scientific literature, and understanding the essential components from research papers.
- Good record keeping practice.
- Hierarchically listing all possible reasons for the failure to obtain results in an experiment with the aim to enhance their trouble shooting abilities.
- To write general Standard Operating Procedures (SOPs) and follow them.
- Handling basic equipment independently.
- Comprehending a given protocol and execute the same independently.

c) Avoiding the usual presumption that students would/should have already learned all essential laboratory practices. Fresh efforts were made to ensure a uniform high-reliability among all students in terms of such basic procedures. The efforts included creating exercises to promote self-realization of the key concepts and self-learning of essential experimental procedures. Improvements in basic operations were also attempted via a combination of pre-performance discussions about simple experiments, instructions during experiments, and post-performance group-analysis of results and potential causes of variations, etc.

d) There was an equal emphasis on teamwork and independent performance. While students were asked to perform several experiments individually, and repeatedly, they also had to frequently work in groups and the group members were often randomly shuffled. The groups were active mostly during the analysis of results.

Discussions to promote self-realization of safety and reliability of results: During the first two months, active sessions were organized to help students realize the significance of safety and record keeping. For example, after an initial narration of a few real and a few hypothetical accidents, their potential causes and preventive measures were discussed. After this, student-groups were asked to state associated hazards and possible preventive measures. They were then prompted to imagine varying circumstances, anticipate/identify possible mishaps and corresponding hazard at every step in a given protocol, and asked to list preventive measures corresponding to each type of potential mishap. Students were also constantly encouraged to admit any mistakes or accidents so that we could all discuss the event and learn from it. It was a fun and learning exercise in most batches, as efforts were taken to ensure that no one feels punished or upset because they admitted spills, or breaking of glassware, etc. Similarly, the significance of correct execution of the routine procedures was frequently discussed, with the intention of inculcating the right kind of basic laboratory habits, particularly during the quantitative experiments.

### Phase II objectives, for the remainder of the in-house training (120-180 days)

Mainly to help the majority of students to acquaint with common molecular biology experiments and develop a confidence in the following aspects.

- To prepare a step-wise molecular biology protocol based on any given paragraph in the methods section of a research paper.
- To identify the apt bio-technique/method for specific purposes.
- To independently design common molecular biology experiments.
- To independently plan, prepare and execute such experiments that require handling DNA, RNA, proteins, recombinant DNA technology, mammalian cell culture and/or plant tissue culture.
- To learn the working of advanced techniques such as microarray, DNA sequencing, 2-D electrophoresis, and FPLC.
- To write correct and unambiguous statements to represent the aim of the experiments and interpretation of the results.

e) Maintaining a good student to instructor ratio. Most of the times, there was one well-trained instructor for a maximum of 5 students. The first batch of 5 students were trained directly by the convener and some of the students from most of the batches were hired as instructors for later batches.

f) Stressing practice of common molecular research methods as well as industrially relevant basic procedures, such as constant and detailed documentation of procedures and observations. [See syllabus in supplementary notes].

g) Employing quantitative experiments repeatedly: In initial batches, students were asked to identify protein concentrations by the Bradford assay. These quantitative experiments took more time. The student-feedback also showed that they did not like putting so much time on one type of experiment. Hence, the titration-based estimations (of glucose, for example) were used in later batches. Each student was asked to repeat a quantitative experiment about 6 times. In most cases, deliberate, specific variations were introduced to provide opportunities to test the influence of procedural variations or change of reagents on the outcome. Students were asked to compare the results, analyze them and perform statistical estimation of the extent of variations in the results of their replicates (across days) as well as of those across class-mates. Frequent sessions were arranged to discuss the statistics after the experiments. Students’ attention was frequently drawn to the need to: (i) follow correct procedures for simple steps such as measurements (volume, weight, pH, etc.), mixing and transferring; (ii) be observant and slow in the beginning to ensure ‘developing the right habits’ in executing such steps; and (iii) develop the right habits early in the program. The difficulty in ‘unlearning the wrong habits’ later (after a few years of experience) was stressed. Mandatory group interactions at different stages allowed them to appreciate a variety of sources of variations in their results, such as the cleanliness of lab-wares, minute changes in basic measurements, brand/lot variations of the chemicals used, and sensitivity and positioning of equipment. Such group discussions often involved analysis of the records of observations. In addition, attempts were made to inculcate the following habits in all students through multiple assessments and verbal reminders: (i) prior work such as writing most of the record-book-entries, discussing the plans but individually preparing for each experiment; (ii) independently recording observations during the experiments; and (iii) group-discussion for a comparative analysis of results. Students were always asked to list possible reasons for variations in the results of each experiment, from electrophoretic analysis of proteins/DNA/RNA samples to cloning experiments, and encouraged to discuss the same with other group members, instructors and the convener. The intension of these efforts was to improve their trouble-shooting ability as well as promote self-learning.

h) There was an equal emphasis on experimental purposes as well as procedures: Every technique/procedure was linked to a specific question where possible. For example, for bacterial growth curve determination, the student-groups were first briefed about the principle of the method and asked to come up with questions for which they would like to find answers through the experiment. They were then asked to design a study to address the selected questions, followed by group-discussions in the presence of the instructors/convener. The groups later proceeded to conduct the experiment. In some cases, a set of related experiments were planned consecutively and thus formed a logical series with specific objectives.

i) Laboratory management: Students had to monitor the usage and functionality, and manage commonly used laboratory equipment. They were assigned complete responsibility of 3 diverse types of equipment by the third month of the program. This included a primary responsibility for one specific equipment and secondary responsibility for two others. The tasks involved were learning the operational and maintenance procedure, preparing an SOP for correct operation, recording the extent of usage and ensuring the constant operational readiness of the equipment. After 2-3 months, the responsibilities were shuffled. Thus, every student experienced maintenance of at least 6 different equipment during the course.

In addition, assignments were given to students to i) familiarize with common manufacturers and distributors of frequently used equipment, reagents and other consumables; ii) compare catalogues; iii) compare quotations and technical features in the context of some of the recent equipment purchases; and iv) periodically monitor the stock of chemicals and other consumables.

## Results

Though the main objectives and contents were constant across 8 batches, the contents were slightly improved almost, and the duration enhanced from 6 to 8 months gradually. We also offered an additional internship for 6 months, as an option, in later batches.

Overall, the results were very satisfying as the majority of students attained the targeted capacities. Most of them, in all 8 batches, seemed to be enabled as expected. Specific results are described below.

### Enhanced precision & accuracy in students’ results

Average variance (mean of absolute difference of each data point from the arithmetic mean of all data points) across triplicates in the results of quantitative experiments improved remarkably during the first or second repetition, in all batches. The initial instructions & the discussions held after the first round of quantitative experiments seem to have helped in bringing such improvements in the results. To test influence of such prior discussions, a special case study was conducted in a batch. Students were randomly grouped into two and asked to repeat protein estimations (Bradford method). While one half of students in the batch were simply told to get correct and reproducible results, the other group received specific tips on ways to reduce variations, before beginning the experiments. These tips and interactive sessions sometimes took about 15 minutes only, even though repeated instructions/reminders helped in case of some students. Examples of tips given included the need to consistently use of one set of (calibrated) measurement equipment for a given experiment, do’s and don’ts when handling cuvettes, pipetting, and transferring, and to keep consistency in each of such procedural details. The positive influence of such instructions was seen across batches. A specific experiment was done once to test this notion. This experiment indicated that simple repetitions, without such special instructions, may not help in improving the reliability of results (figure 1) - even though the students felt confident in getting reliable results by just repeating the experiments. Reductions in variations in volumes of solution consumed during titration experiments were consistent despite introduction of a new type of procedural (figure 1) variation such as measuring Cu & Ni by titration-methods, instead of glucose estimations with which they started the titration experiments. In a separate experiment, it was observed that significantly (P <0.0001, paired student t-tests) higher precision (mean variance across triplicates) and accuracy (correlation between spectrophotometric readings and protein concentrations) was noticed in a group (n=20) that received the special instructions listed above, than the control group (n=19) that received no such instructions.

**Figure 1.**
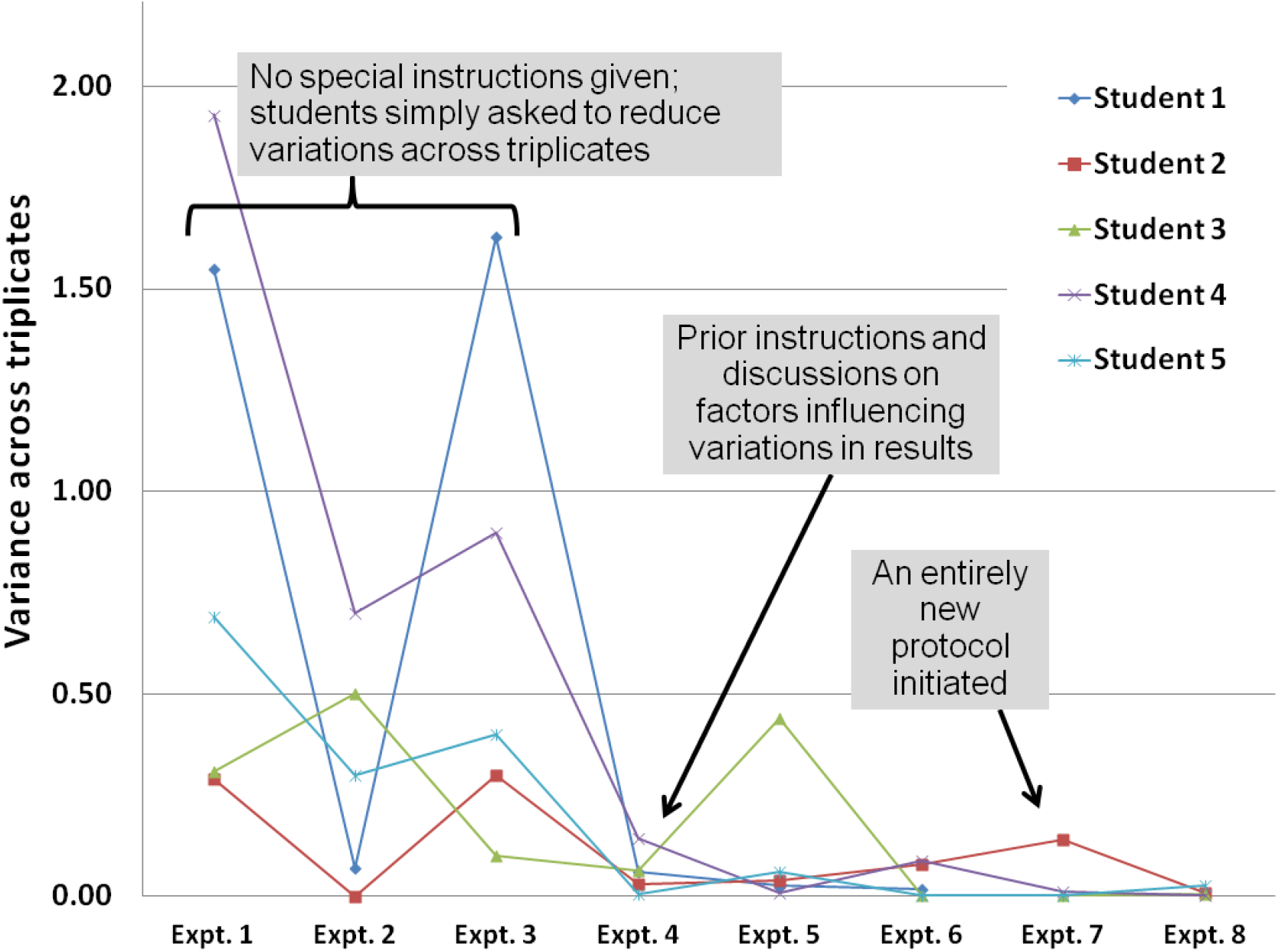
Variations in results during titration-based experiments in a group of LCBT students of one of the batches. The first 3 experiments were performed without any tips while the next 3 were performed after intervening instructions and discussions on ways to reduce variations. Variance in the volume of sodium thiosulphate used during titration experiments for estimating glucose in a given solution is depicted in the graph for experiments 1 to 6. A completely new titration protocol (to estimate copper in a solution) was given to students and variance was again measured by them in experiments 7 & 8. Student 4 did not perform the last 2 experiments. NOTE: While such experiments could not be done by involving a higher number of students, it should be noted such studies were conducted to test if a trend noticed informally across multiple batches of students.

### High success rate

About 65% of students across 8 batches scored more than 70% marks [an example of students’ performance in Table S1]. Team-work culture, individual dependability, and practices in documentation and laboratory management were imparted well among most of the students. More than 80% of students in each batch attained proficiency in basic skills and habits required for general research and good laboratory/manufacturing practices (GLPs/ GMPs), such as creating and following SOPs, solution preparations, prompt record-keeping of plans, procedures, observations and results, and reliable execution of protocols. By the third month of the course, almost every student was capable of independently performing a given new task without prior demonstration or explanation. For example, about 90% of the students could obtain expected results in their first attempt of the extraction of total DNA, proteins, and RNA. Similarly, they could also analyze the quality of such macromolecules extracted by them via the standard electrophoretic and spectrophotometric analysis, and quantify the same.

### Evaluations were continuous

Record book monitoring, assessment of the execution of procedures, adherence to SOPs, responsible roles in equipment maintenance, completion of an assignment, quality of results, etc., were routinely and independently assessed by the instructors and a grade was assigned based on the average scores across instructors. Besides, two formal practical tests were conducted to assess different aspects. A cumulative score out of 100 was used to derive grades for each student at the end of the program, as indicated here: D: below 31, C: 31-50, B: 51-60, B+: 61-70, A: 71-80, A+:81-90, & S: 91-100. Across batches, no student scored an overall ‘D’ (lowest) OR ‘S’ (highest) grade. About 65 of the total 101 students from 8 batches obtained an overall ‘A’ or ‘A+’ grade, and 5 students ended up with an overall ‘C’ grade. The students were graded under the following specific categories: A. General skills related to professions in biotechnology (computer skills, basic bioinformatics, mathematics for biology and basic statistics; concepts in research, molecular biology (theory), literature search and presentation skills; general laboratory discipline, maintenance and safety aspects; designing experiments, interpreting results and record-keeping); B. Accuracy and precision of results; C. Basic procedures in handling nucleic acids (extraction and quantification; analysis by gel electrophoresis; primer designing, PCR, RT-PCR, DNA, and RNA-blotting procedure); D. Recombinant DNA procedures (restriction digestion, ligation, preparing competent cells, transformation, expression of cloned sequences and cell culture); E. Basic procedures in handling proteins (extraction and quantification; protein purification by chromatography and other techniques; Western and ELISA; native and SDS-PAGE; staining procedures); F. Internship/project. The average percentage of students who scored S, A+, A, B+, B, C, and D grade in any specific category, across all batches, was 10, 18, 42, 15, 8, 6 & 1, respectively. Thus, nearly 70% of students scored more than 70% in most categories. Even though a complete study could not be done to analyze the results, students from various types of economic, geographical, prior-studies and social backgrounds seemed to perform equally well.

Along with independent functioning, most students were also comfortable with team-work. Laboratory accidents such as spillage or breaking of glassware were rare - especially after the first month of the program. All students also successfully managed utility and log-books of the laboratory equipment and consumable stocks. The instructors had less role to play in general laboratory management after the first 3-4 months. Parts of discussions towards improved safety, accuracy and precision not only seem to have helped in enhancing these parameters in later experiments, but they may also have contributed in the creation of a better SOP for specific applications. Such SOPs, created for one specific purpose in each batch, were then compared with the authentic SOPs prepared by an expert. Students also enjoyed active group discussions for deciding the purchase of the right laboratory reagents/equipment required for IBAB. For example, real-time PCR equipment was purchased following a debate across 3 student-groups of a batch. These students studied multiple models available in the market in terms of technical features as well as cost.

Almost all experiments taught in the program were not performed earlier by most students, independently at least. Interestingly, the majority of the course participants also learned many of the following basic aspects for the first time during the current program: advanced computer operations with spreadsheets and presentations, moderately advance handling of text documents, essential laboratory safety procedures and precautions, micro-pipetting, literature search, basic statistics, and the fundamental concepts in IPR, GLPs, and GMPs. Besides, almost all students were used to the practice of making the required entries in their laboratory records long after the actual experiments. During LCBT, however, it was mandatory to enter the requirements, the protocol and tables for observations in the record books, before the actual performance of the experiments. The observations and data entries were made during the experiment. Only result-interpretations were allowed to be written after the experiments. All students developed this as a habit and most of them found these record keeping habits useful for various purposes, including trouble-shooting. The efficiency in making neat entries increased with time in most of the students in each batch. It was also compulsory for each student to list potential reasons for any failure or deviations from the expected results and hierarchically arrange these reasons in the order of probability, and discuss the same with instructors and/or class mates.

### Placements

Although it was a challenge for the convener of the program (author) to place the students, particularly due to small batches and newness of the program, the novel student profiles (S1 table is an example) received a positive response and eventually placements occurred at a reasonable rate. In many cases the students were first recruited as interns by Companies, and more than 85% of the students placed as interns received a stipend. The internships seemed to boost their confidence and help them learn more in specific areas of research or industrial applications. It also gave the recruiting organizers a chance to test the students over 6 months before hiring. In most cases, students were hired by the host organizations soon after their internship. A few internship-supervisors from different companies voluntarily gave positive feedbacks about the LCBT students recruited by them.

Even though the initial batches were small a total of 102 students completed the LCBT across 8 batches. Of these outgoing students a few opted for further studies and about 80 started their professional life, usually within the first month of course completion (see table 1 for details). Many organizations, particularly smaller and younger companies, which recruited our students contacted us later to express their continued interest in hiring the next batch of students. Thus, placements became much easier after the initial batches. In fact, during later batches the placements received a stiff competition due to the advent of another nation-wide program called Biotechnology Industrial Training Program (see below), where the industry received financial benefits for training new graduates/post-graduates, who also received a stipend. Despite this, however, LCBT students were preferentially recruited, particularly by established bigger organizations. Even though confidence and real capacities were developed among students during the program, not having a formal master’s degree was perceived as a set back by some of the students. Hence, a small portion of students took to other higher studies such as MTech or MSc, immediately after LCBT. The immediate placement record of each batch of each student of this program, just like the other contemporary programs at IBAB including the diploma in bioinformatics, were made available on the institute’s website.

**Table 1.** Statistics of admissions and placements across LCBT batches

| Batch (& year) | No. of applicants | Total no. of students enrolled | No. of students who discontinued LCBT | No. of students who took a job in different areas* |  |  |  |
| --- | --- | --- | --- | --- | --- | --- | --- |
|  |  |  |  | No. of students pursued further studies* | Academic research | Industrial position | Teaching |
| 1 (2004) | 29 | 5 | 1 | 0 | 1 | 0 | 3 |
| 2 (2005) | 130 | 11 | 0 | 0 | 4 | 4 | 1 |
| 3 (2006) | 66 | 9 | 1 | 1 | 1 | 2 | 1 |
| 4 (2007) | 128 | 19 | 1 | 1 | 1 | 15 | 1 |
| 5 (2008) | 118 | 12 | 0 | 2 | 0 | 2 | 4 |
| 6 (2009) | 135 | 17 | 0 | 0 | 7 | 5 | 4 |
| 7 (2010) | 132 | 14 | 0 | 1 | 2 | 8 | 0 |
| 8 (2011) | 121 | 18 | 0 | 4 | 7 | 3 | 3 |
| Total | **859** | **105** | **3** | **9** | **23** | **39** | **17** |
|  |  |  |  | **88** |  |  |  |

\*A few students did not keep in touch with the convener &/or did not find employment even after 2 or more months after completion of the program.

Most of the alumni members seem to now have a steady career in various life sciences domains. A few more (about 13) ex-students completed Ph.D. in India or abroad, and are currently engaged in high-quality research as independent scientists at various organizations in India and abroad. At least three alumni have tried entrepreneurship (2 companies started between them). A few have grown into allied careers such as intellectual property rights management and regulatory affairs. Most others have been assisting R&D activities in different Indian companies.

Overall, eight batches of LCBT suggest that the novel combination of approaches tried here could be used to a) meet the challenge of dealing with the highly divergent initial capacities of students in various aspects, in developing countries such as India; b) build up a reliable technical workforce for the biotech industry, and c) prepare students for academic research, involving laboratory work in cell and molecular biology.

## Discussion

The objectives of the course were achieved and the program was generally successful in developing the capacities originally planned for. But some of the ambitious training-goals could not be achieved. For example, in their first attempt, about 30% of students could not amplify cDNA (RT-PCR) corresponding to a specific mRNA using the RNA isolated by them individually; they were successful only in their second or third attempt. In fact, only about 20% of students succeeded in correctly executing all steps for cloning the assigned genes, which formed the last assignment (with a time limit of 2-3 weeks) of the course. The program could also not focus on improving the originality of scientific thinking, theoretical knowledge of related subjects and the context of techniques. In some batches, for reasons not understood well, students hesitated to come forward admit procedural mishaps that could be discussed to learn more about possible preventive measures, etc. Although active learning strategies (12–16) were employed, time limitations did not permit engaging students in inquiry-based learning to the required extent. For example, the detailed discussions on the questions or the context of inquiry for the growth curve experiment was indeed rare. The discussions were kept very brief for most cases. Of course, such goals of integrated teaching of multiple facets could be a challenge even in programs of longer duration (17).

Apart from the time constraints, there could be a few other reasons for the observed limitations. Instructors’ growth and maturity was crucial to the program. While most instructors put in great efforts, constant change of instructors created a severe set-back. It was also a learning process for the convener (author). Convening all 8 batches and being actively involved in instruction and group discussions, as well as continuously seeking feedbacks, did help the convener learn and improve the program. About 100 hours dedicated every year by him for teaching, discussions, etc, was perhaps not enough. Apart from carrying out his own research, he had a few other parallel responsibilities at the institute, including contributing to the teaching (about 100 hours a year again) in various topics as part of a separate bioinformatics post-graduate diploma course.

The Indian education system has faced several challenges and has undergone many improvements over time (18, 19). I hope the reported humble efforts trigger many more improvements within the life sciences domain. In the current program, the convener planned the sessions and closely monitored the progress across the days in each batch, which were not large. But, it may be prudent to use well-trained instructors (see 20). It may also be important to monitor and guide the discussions among students and have followed up feedback sessions rather than having student only discussions (see 21).

There have been other efforts in India to address the current shortcomings. For example, a Biotechnology Consortium of India Limited (BCIL), an organization promoted by the Department of. Biotechnology, Government of India, has initiated a Biotech Industrial Training Program (BITP) (http://bcil.nic.in/biotech_industrial-training.html) since several years. Under this scheme, selected youngsters, with a recent master’s biotechnology-related degree, can undertake training directly in the private sector while the BCIL pays both the trainee and training organization. Similarly, a BioTechnology Finishing School (BTFS) has been initiated by the Department of IT, BT and S&T, Government of Karnataka, and supported by the Department of Biotechnology, Ministry of Science & Technology, Government of India. BTFS has been recently replaced with a similar new program named BiSEP (http://bisep.karnataka.gov.in/). Under these schemes, selected educational centers are supported for specialized training for about 6 months. The students receive a stipend from the government while they undergo such training. While such efforts have their own merit in addressing the human resource development needs, there is an urgent need to enhance the basic laboratory training as well as teaching to impart the fundamental concepts in regular bachelors’ degree courses as well as earlier educational programs. The lessons learned from LCBT could help in developing better training modules as part of the regular bachelor’s and/or master’s programs or as an alternative diploma after a bachelor’s degree in any life sciences stream.

## Conclusion

The consistent placement record, and the follow-up growth in the career of most alumni members, indicates that the program was largely successful in achieving the objectives. In every batch, the efficiency and dependability of most students constantly increased during the course. Complete freedom and support were given by the institute, particularly in the initial years, and a personal passion for teaching contributed to achieving the goal.

The current proof of the success of LCBT and insights into the training strategy could be useful in framing novel courses and curricula in life sciences programs in developing countries, particularly in places where a highly diluted talent-pool is generated in the life sciences sector every year. The LCBT structure could be modified, improved and adapted for different purposes such as exclusive research methodology training or building technical work-force for general industrial purpose and/or a specific type of industry.

## Acknowledgments

The institute has been initiated, and generously supported by KBITS, ITBT dept., Government of Karnataka. It has also been supported directly or indirectly by multiple departments of the Government of India, such as DEITY and DST-FIST.

## Supporting information

### Supplementary note 1. Selection Process

Applicants with a bachelor’s or higher degree in any life sciences stream with a sound theoretical cell and molecular biology knowledge were invited to take an online entrance test (conducted by Shodhaka: www.shodhaka.com/SOTS) which comprised of multiple-choice questions related to general aptitude, English, and fundamentals of Biology, Chemistry, Physics and Mathematics. The test was a means to screen candidates eligible for an interview and could be taken from any place with good computer and internet facilities. A 40 to 50% scoring limit was used to short list applicants for interview. The interviews were conducted by a panel of at least three scientists led by the convener. Purpose of the interview, possible areas of discussions, thresholds for qualifying, the need to ensure that the candidate is not nervous, etc. were discussed before beginning the interviews that lasted about 45 minutes per candidate across 2-4 days every year. All panel members scored the candidates individually on a 1 to 100 scale. It was pre-decided that in most cases anyone scoring below 50 would not be offered admissions to the course. Although, up to 25 students could be selected, we could only select lesser candidates as most did not perform well, which in turn was because they did not have the required clarity about the fundamental concepts in cell and molecular biology, without which the hands-on training would not make much sense. Overall, about 1 in 6 applicants was offered admission to the program.

### Supplementary note 2. The syllabus

details are given below. The curriculum covered most of the commonly employed molecular biology experiments as well as the basic and general laboratory practices. The specific contents were continuously modified to improve the quality of training across batches. For example, earlier batches did not have a few components. But most of the yearly changes were in terms of the time spent on each component. Several other basic skills, including English writing and computer operations (see objectives in the main text) were considered as key objectives based on observations during early batches. In all batches, most of the in-house training involved planning, preparing, performing independent experiments by the students and/or discussions on all aspects including variations in observations and result-interpretations. Such opportunities were created as much as possible with an intention of promoting self-realization and self-learning among the students. After the initial training, during the last six months of the program, selected students also undertook an individual basic-research or industrial project work.

The theoretical understanding of many students were found to deteriorate in first 3 batches, as observed during intermittent informal discussions and mock-interviews. Hence, periodical lectures and discussions on fundamental cell and molecular biology concepts were included in the later batches. Students were encouraged to actively participate in guest lectures on advanced research topics, which were organized at regular intervals at the institute, to help them recollect the theoretical aspects, and inspire them for advanced readings.

#### A. Theory with practical sessions, discussions or assignments

- Cell and molecular biology: A brief refresher course
- Research in life sciences: An overview and a few specific examples.
- Safety, common equipment and reagents used; basic computer skills and statistics.
- Introduction to bioethics, IPR issues, some of industrial activities and practices and related statutory regulations.
- Record keeping (patent-oriented) and other relevant professional skills (e.g., presentation skills, literature search, experimental designs, interpretation of results, and team work).
- Understanding accuracy, precision and variations in results.
- Basic bioinformatics: Sequence analysis, primer designing, literature search and related databases and tools.

#### B. Practical sessions with multiple (repeated 2-5 times each) individual performances by the students

- Basic bacterial culture methods.
- DNA and RNA handling: genomic DNA (plant, microbial and animal) isolation and quantification.
- Analysis of DNA samples by gel electrophoresis.
- RNA isolation.
- Quantification and gel electrophoresis of RNA.
- Recombinant DNA technology.
- Primer designing and PCR.
- Restriction digestion, ligation, competent cells preparation and transformation.
- Cloning and plasmid isolation.
- Protein handling - Extraction and quantification, Native and SDS-PAGE; staining procedures for the same.

#### C. Practical sessions with limited number (1-3 times each) of individual performances by the students

- Purification of DNA after enzymatic reactions
- mRNA isolation
- Reverse transcription-PCR
- Blotting DNA and RNA for Southern and Northern
- Two-dimensional electrophoresis
- Expression of cloned sequences
- Western and ELISA
- Protein purification: salt precipitation and chromatography
- Mammalian cell culture

#### D. Demonstrations only

- Microarrays
- DNA sequencing
- Protein activity assays
- HPLC

#### E. Second semester (6 months)

*Internship (mostly with stipend), of six months in a life sciences organization, for selected students.*

**Table S1:** An example of students’ profile (7^th^ batch) used for placement efforts towards the end of the program. The final transcript used grades of D, C, B, B+, A, A+ & S (corresponding to 0% to 30%, 31 to 50%, 51 to 60%, 61 to 70%, 71 to 80%, 81 to 90% and 91 to 100% respectively).

| Name of the student<br>(actual names not shown here as the purpose is show an example of the profile of students' scoring, a profile that was used for seeking placement for students) | Previous degree and the main subject<br>(*biotech* = *biotechnology*; *micro* = *microbiology*) |  | General skills related to professions in biotechnology: Computer skills, basic bioinformatics, mathematics for lab-biology and basics statistics, literature search, presentation skills, record keeping & lab-maintenance. | Accuracy and precision of results | DNA - handling | RNA handling | Protein handling | Recombinant DNA technology, mammalian cell culture etc | Overall practical abilities | Theory: cell & molecular biology, & principles of techniques | Overall score (%) | Current placement status (recruiting organizations are indicated for those placed already) |
| --- | --- | --- | --- | --- | --- | --- | --- | --- | --- | --- | --- | --- |
| Student 1 | BE | Biotech | 80 | 81 | 76 | 68 | 72 | 86 | 464 | 81 | 78 | Institute of micronutrient technology, Bangaluru |
| Student 2 | BE | Biotech | 88 | 77 | 66 | 66 | 72 | 76 | 445 | 95 | 77 | Indian institute of science, Bangaluru |
| Student 3 | MSc | Biotech | 89 | 72 | 71 | 57 | 76 | 63 | 429 | 90 | 74 | Shodhaka LS pvt ltd, Bangaluru |
| Student 4 | BE | Biotech | 93 | 82 | 75 | 69 | 79 | 79 | 477 | 92 | 81 |  |
| Student 5 | BTech | Biotech | 81 | 70 | 66 | 55 | 77 | 38 | 387 | 65 | 65 |  |
| Student 6 | BE | Biotech | 85 | 75 | 70 | 65 | 70 | 53 | 417 | 89 | 72 | La ReUnion, Internships |
| Student 7 | BE | Biotech | 77 | 77 | 67 | 56 | 71 | 84 | 431 | 71 | 72 | Pursuing higher studies |
| Student 8 | BTech | Biotech | 81 | 74 | 66 | 63 | 73 | 56 | 413 | 83 | 71 |  |
| Student 9 | BTech | Biotech | 74 | 76 | 65 | 63 | 68 | 68 | 412 | 67 | 68 | Agilent technologies, Bangaluru |
| Student 10 | MSc | Biotech | 72 | 61 | 69 | 52 | 68 | 53 | 375 | 83 | 65 | LOOKING FOR A JOB / paid internship |
| Student 11 | BE | Biotech | 84 | 67 | 67 | 55 | 69 | 38 | 381 | 69 | 64 |  |
| Student 12 | BTech | Biotech | 76 | 68 | 55 | 67 | 72 | 38 | 376 | 75 | 64 | Novozymes, Bangaluru |
| Student 13 | BE | Biotech | 69 | 80 | 75 | 62 | 76 | 87 | 449 | 76 | 75 |  |
| Student 14 | MSc | Micro | 57 | 50 | 69 | 40 | 65 | 36 | 317 | 86 | 58 | Biozeen, Bangaluru |

These records were about 10 days before the completion of the specific batch. In most batches the placement process for most students was completed before the course completion. Difficulties were faced in placing students when some of them insisted very specific domains (such as academic position in specific cities)

